# A FOXO-dependent replication checkpoint restricts proliferation of damaged cells

**DOI:** 10.1101/2020.01.09.900225

**Authors:** Marten Hornsveld, Femke M Feringa, Lenno Krenning, Jeroen van den Berg, Lydia MM Smits, Nguyen BT Nguyen, Maria J Rodriguez-Colman, Tobias B Dansen, Rene H Medema, Boudewijn MT Burgering

**Affiliations:** Oncode Institute, Department of Cell and Chemical Biology, Leiden University Medical Center, 2333CZ, Leiden, The Netherlands; Oncode Institute, Center for Molecular Medicine, University Medical Center Utrecht, Utrecht University, 3584CG, Utrecht, The Netherlands; Oncode Institute, Division of Cell Biology, Netherlands Cancer Institute, 1066CX, Amsterdam, The Netherlands; Department of Molecular and Cellular Neurobiology, Faculty of Science, Center for Neurogenomics and Cognitive Research, Amsterdam Neuroscience, Vrije Universiteit Amsterdam, 1081HV, Amsterdam, The Netherlands; Oncode Institute, Hubrecht Institute–KNAW (Royal Netherlands Academy of Arts and Sciences), 3584CT, Utrecht, The Netherlands

**Author notes:** These authors contributed equally to this work (Marten Hornsveld, Femke M Feringa). Correspondence can be send to Marten Hornsveld and Boudewijn MT Burgering, and.

## Abstract

DNA replication is challenged by numerous exogenous and endogenous factors that can interfere with the progression of replication forks. Stalling or slowing of the replication fork as a result of replication stress leads to formation of aberrant single-stranded DNA (ssDNA) stretches and potentially DNA double-stranded-breaks (DSBs). Accumulation of ssDNA activates the ATR-dependent DNA replication stress checkpoint response that slows progression from S/G2- to M-phase to protect genomic integrity (1). However, whether mild replication stress restricts proliferation remains controversial (2–6). Here we identify a novel cell cycle exit mechanism, that prevents S/G2 phase arrested cells from undergoing mitosis after exposure to mild replication stress through premature activation of the CDH1 bound Anaphase Promoting Complex / Cyclosome (APC/C^CDH1^). We find that replication stress causes a gradual decrease of the levels of the APC/C^CDH1^ inhibitor EMI1/FBXO5 through Forkhead Box O (FOXOs) mediated repression of its transcriptional regulator E2F1. By doing so, FOXOs limit the time during which the replication stress checkpoint is reversible, and thereby play an important role in maintaining genomic stability.

## Results

Mild replication stress leads to under-replicated DNA which may be tolerated by cells, even though this gives rise to DNA lesions upon mitotic progression (3,4). Indeed, replication stress may induce gaps, breaks and micro-deletions at common fragile sites (CFS)(7,8), highlighting its mutagenic potential. Therefore, it is likely that cells have mechanisms to prevent the propagation of under replicated DNA. To investigate this, we visualized the cellular response to mild replication stress by aphidicolin in non-transformed RPE-1 cells with endogenously tagged Cyclin B1^YFP^ (RPE-*CCNB*^YFP^) and stable expression of *53BP1*^mCherry^ (RPE *CCNB*^YFP^-*53BP1*^mCherry^)(9) (Fig. 1A). While the addition of aphidicolin had no effect on cells in G2-phase (identified by Cyclin B1 expression), cells that progressed through S-phase in the presence of aphidicolin showed a clear decrease in mitotic entry over time (Fig. 1B). This decrease in mitotic entry was accompanied by a concomitant increase in cells that abruptly lost Cyclin B1 expression (Fig. 1C). Based on previous findings (10) we hypothesized that the loss of Cyclin B1 expression may correspond to levels of DNA damage. Indeed, cells that degrade Cyclin B1 after progressing through S-phase with mild replication stress, display increased 53BP1 foci compared to control cells or cells that recovered from replication stress and progressed to mitosis (Fig. 1D). Importantly, abrogation of DNA damage signaling by ATR inhibition caused an increase in 53BP1 foci number in G1 daughter cells (Fig. S1A).

**Figure 1:**
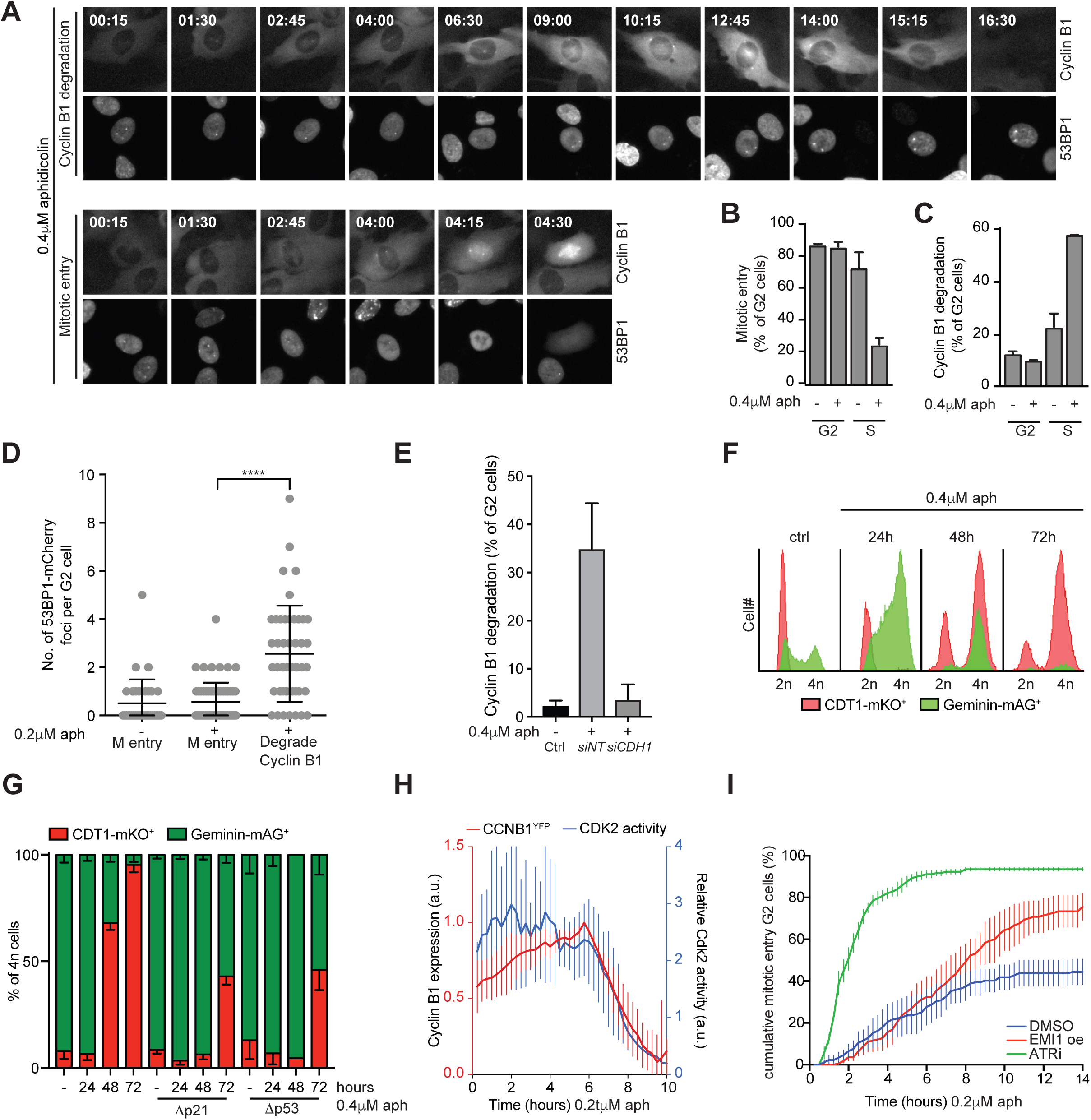
Replication stress leads to APC^CDH1^ activation in G2 phase. **A** Representative image from time-lapse movies of RPE CCNB^YFP^–53BP1^mCherry^ cells treated with 0,4µM aphidicolin. **B** Mitotic entry of G2 cells (Cyclin B1 positive at t=0) or S phase cells (Cyclin B1 positive at t=15) in the presence or absence of 0,4µM aphidicolin. Average of three independent experiments + sem. **C** Percentage of cells from figure B that degrade Cyclin B1 in the presence or absence of 0,4µM aphidicolin. Average of three independent experiments + sem. **D** Quantification of 53BP1mCherry foci in G2 cells before onset of Cyclin B1 degradation or mitotic entry. Dots represent individual cells pooled from two independent experiments. Mean + sd. **** P<0.0001 (Welch’s corrected unpaired t-test). **E** Percentage of G2 cells that degrade cyclin following aphidicolin treatment in presence of *FZR1* knock-down. **F** Representative flow cytometric analysis of hoechst stained RPE-FUCCI cells after 0,4μM aphidicolin treatment for 24, 48 and 72 hours. Red: CDT1-mKO+ (G1-phase), green: Geminin-mAG+ (S/G2/M-phase). **G** Flow cytometry quantification of cell cycle distribution of 4n cells in hoechst stained RPE-FUCCI cells knockout for p53 (Δp53) and (Δp21) (n=3). Red: CDT1-mKO+ (G1-phase), green: Geminin-mAG+ (S/G2/M-phase). **H** Relative CDK2 activity and cyclin B1^YFP^ intensity in individual G2. Lines represent average of 16 cells pooled from two independent experiments + 95% CI. **I** Cumulative mitotic entry of G2 cells that progressed through S phase after treatment with 0,2µM aphidicolin in presence or absence of EMI1 overexpression or ATR inhibitor. Average of three independent experiments + sem.

We and others have previously shown that loss of Cyclin B1 and other cell cycle targets after DNA double-stranded breaks (DSBs) is mediated via APC/C^CDH1^-dependent protein-degradation, causing the cells to enter a state of senescence (10–12). To examine whether the aphidicolin-induced loss of Cyclin B1 is caused by APC/C^CDH1^ activation, we depleted CDH1. Indeed, CDH1 knock-down prevented Cyclin B1 degradation in cells that progressed through S-phase in the presence of aphidicolin (Fig. 1E & S1B). Experiments using RPE-1 cells stably expressing the Fluorescent Ubiquitination-based Cell Cycle Indicator (FUCCI) probes hCDT1 (aa30/120)-mKO2 (degraded by SCF^SKP2^ in S/G2/M-phases, thus indicating G1-phase) and hGeminin(aa1-110)-mAG (degraded by APC/C^CDH1^ in G1-phase, thus indicating S/G2/M-phases) confirmed that APC/C^CDH1^ activation indeed occured in G2. Aphidicolin treatment of RPE-FUCCI cells resulted in the accumulation of 4n mKO2^+^ cells over time, showing that these cells lost the APC/C^CDH1^ target Geminin and SCF^SKP2^ activity without going through mitosis (Fig. 1F). We and others previously demonstrated that the balance between a reversible arrest and the APC/C^CDH1^-dependent cell cycle exit caused by DNA double strand breaks in G2-phase is be tipped towards cell cycle exit by p53-dependent p21 expression (10,11). Interestingly, we observe that APC/C^CDH1^ activation in response to aphidicolin is delayed but not abrogated in p21 (Δp21) and p53 (Δp53) knockout RPE-FUCCI cells (Fig. 1G).

In G2, activation of APC^CDH1^ is prevented by CDK1/2-dependent phosphorylation of CDH1 (13,14). To characterize CDK activity following replication stress, we used the previously described CDK2-activity sensor in RPE *CCNB*^YFP^ cells (9,15). We find that CDK2 activity is stable in G2 cells following mild replication stress in S phase and drops in synchrony with Cyclin B1 (Fig. 1H). The simultaneous loss of CDK2 activity and APC/C^CDH1^ target Cyclin B1 is surprising, since APC/C^CDH1^ activation in response to DNA double strand breaks is described to depend on the premature loss of CDK activity (10,11,16–19). Therefore we expected CDK2 activity to drop before Cyclin B1, yet our results implicate that APC/C^CDH1^ can be activated despite the present level of CDK2 activity.

In addition to CDK1/2 mediated inhibition, the APC/C^CDH1^ inhibitor EMI1 blocks APC/C^CDH1^ activity during S/G2 and loss of EMI1 is sufficient for premature activation of APC/C^CDH1^ in G2 (17,20–23). We therefore combined our RPE *CCNB*^YFP^-*53BP1*^mCherry^ cells with doxycycline inducible EMI1-overexpression (9), to define the importance of EMI1 loss for APC/C^CDH1^ activation following replication stress. Overexpression of EMI1 prevented Cyclin B1 degradation and rescued mitotic entry of cells exposed to replication stress (Fig. 1I & S1C). As replication stress results predominantly in ATR/CHK1-activating lesions (24), this raises the question whether sustained ATR-signaling is required to maintain cells arrested and induce cell cycle exit. To address this, we treated G2 cells that progressed through S phase in the presence of aphidicolin with the specific ATR inhibitor VE-821. ATR inhibition prevented Cyclin B1 degradation and resulted in immediate mitotic entry of almost all G2 cells that progressed through S phase in presence of aphidicolin (Fig. 1I & S1C). In contrast, aphidicolin-treated cells that did recover from replication stress in the DMSO-treated condition showed a clear delay in mitotic entry (Fig. 1I). The rescue by ATR inhibition was distinct from the rescue observed upon EMI1 overexpression, since EMI1 expressing G2 cells still arrested for several hours before progressing into mitosis. These results show that sustained ATR activity maintains the G2 arrest that ultimately results in APC/C^CDH1^-dependent cell cycle exit. Together, these results imply there are additional players in the replication stress checkpoint that steer the decision between entering mitosis or exiting G2 through regulation of APC/C^CDH1^ activation. Surprisingly, this regulation is not mediated through p53/p21 or Cyclin B/CDK1 signaling.

DNA damage results in the activation of a transcriptional program that promotes DNA repair and stalls cell cycle progression (25). To determine whether a similar transcriptional program is induced by the canonical replication stress checkpoint, we determined the transcriptome of control and aphidicolin-treated cells. We identified 272 genes that were differentially expressed upon replication stress (Fig. 2A, log2fold - 0.5<x>0.5, p <0,05, 155 down- and 117 up-regulated genes). Transcription factor binding sites (TFBS) analysis revealed a significant enrichment of Forkhead Box O (FOXO) target genes that were upregulated in response to aphidicolin (Fig. 2A-B). As FOXOs are known regulators of cell cycle arrest and are involved in the DNA damage response (26,27), we determined if FOXOs are activated in response to replication stress. Indeed, we observed an increase in nuclear FOXO3, specifically in S/G2-phase cells already at 2 hours after aphidicolin treatment (Fig. 2C,D). Although FOXOs usually function redundantly and mRNA expression of *FOXO*1, *FOXO3* and *FOXO4* increases after aphidicolin treatment, FOXO4 protein is undetectable in RPE cells (28) and FOXO1 localization was hardly affected by aphidicolin treatment, suggesting a more dominant role for FOXO3 in response to replication stress (Fig. S2A,B). Nuclear translocation of FOXO3 was reduced in the presence of ATR inhibitor, showing that ATR somehow influences FOXO3 activation in response to replication stress (Fig. 2D).

**Figure 2:**
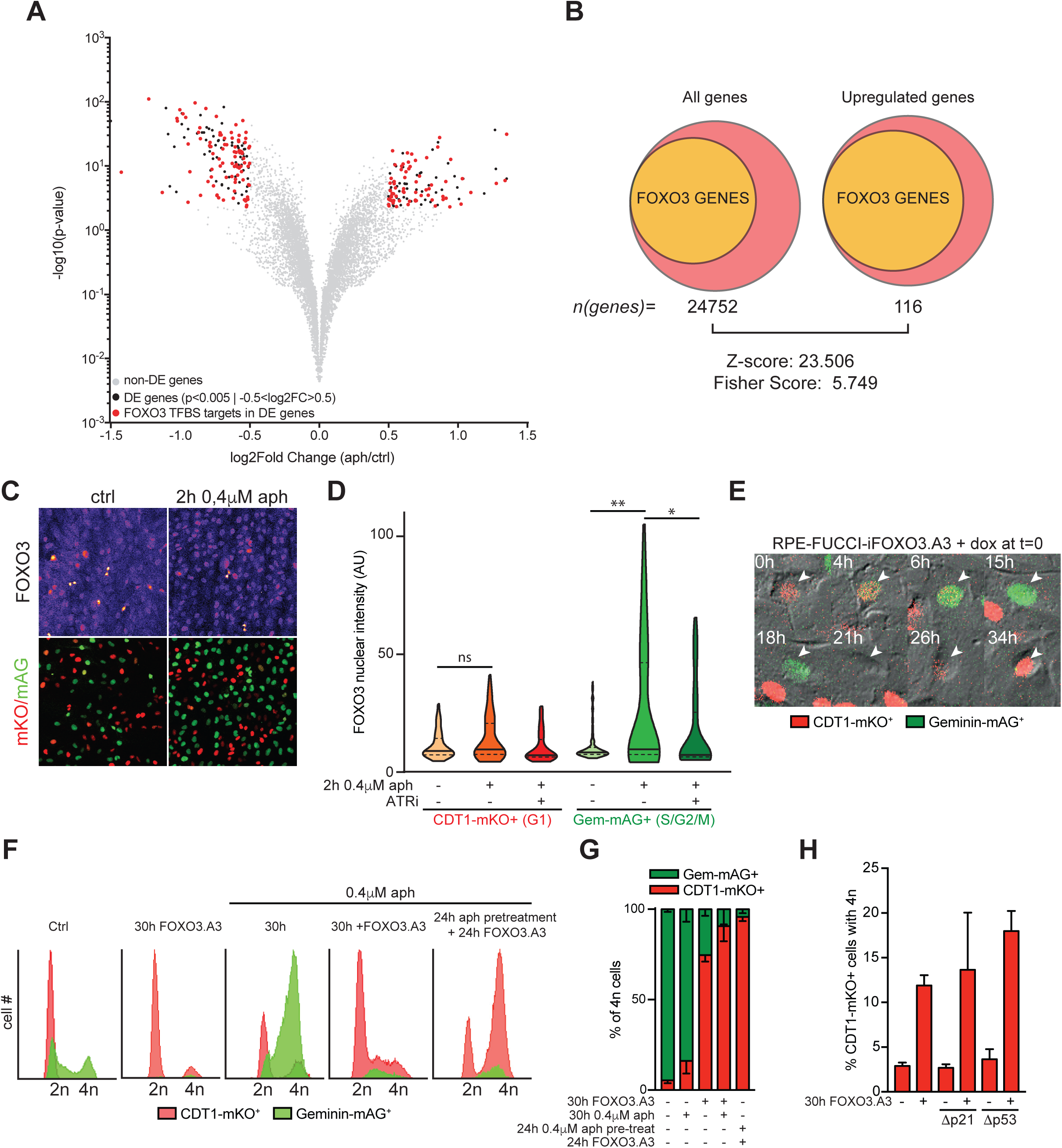
FOXO3 is activated by replication stress and induces cell cycle exit from G2 phase. **A** Volcano plot showing differentially expressed genes comparing aphidicolin-treated RPE-1 cells (7hrs) vs untreated RPE-1 (non DE = grey, DE = Black ((p<0.005 | - 0.5<log2FC>0.5)), DE genes with at least 1 FOXO Transcription factor binding sites (TFBS) = red) after RNA-sequencing of aphidicolin treated RPE cells. **B** Venn diagram illustrating the overrepresentation of FOXO TFBS in promoters amongst upregulated genes (116) to the total all promoters (24752). **C** Immunofluorescence image of FOXO3 (Grey) expression and localization in RPE-FUCCI cells. Red: CDT1-mKO+ (G1-phase), green: Geminin-mAG+ (S/G2/M-phase). **D** Violin plot of average nuclear intensity of FOXO3 in the absence/presence of 0,4μM aphidicolin or 5μM ATR inhibitor VE-821. Solid line = median, dotted line = quartile. Welch’s corrected unpaired t-test P<0.05=*, p<0.05=**. **E** Representative fluorescence time lapse images of RPE-FUCCI-iFOXO3.A3 cells that shows cell cycle exit 18h after FOXO3.A3 induction. A geminin-mAG (green) positive cell becomes CDT1-mKO (red) positive without progressing through mitosis. **F-G** Flow cytometric analysis example and quantification of hoechst stained RPE-FUCCI cells after 30h FOXO3.A3 expression (FOXO ON), 30h 0,4μM aphidicolin, combined treatment and FOXO3.A3 expression in cells pretreated with aphidicolin for 24h. Red: CDT1-mKO+ (G1-phase), green: Geminin-mAG+ (S/G2/M-phase). **H** Flow cytometry quantification of 4n CDT1-mKO+ cells after FOXO activation in p53 (Δp53) and p21 (Δp21) knockout RPE-FUCCI cells (n=3).

To study whether nuclear localization of FOXO is sufficient to cause premature APC/C^CDH1^ activation, we transduced a doxycycline-inducible, constitutively nuclear mutant of FOXO3 (FOXO3.A3) in our RPE-FUCCI cells (Fig. S2C), and analyzed the fate of cells with activated FOXO3 by time lapse microscopy. FOXOs are known to induce a robust G1 cell cycle arrest, and therefore only cells in early S/G2 at the time of FOXO3.A3 induction were included in our analysis (29). Upon FOXO3.A3 expression cells arrest for 15-35 hours in S/G2-phase before prematurely activating the APC/C^CDH1^, much alike the cellular response to replication stress (Fig. 2E). Additionally, we find that 74±4% of all 4n cells lost geminin expression (mAG-) and are mKO+ at 30h after doxycycline-induced FOXO3.A3 (Fig. 2F-G), indicative of APC/C^CDH1^ activation and cell cycle exit from G2 without mitosis. FOXO3.A3 expression strongly enhances aphidicolin-induced premature APC/C^CDH1^ activation, both when FOXO3.A3 is induced simultaneously with aphidicolin treatment or following 24h pre-treatment with aphidicolin, which accumulates cells in G2 prior to FOXO3.A3 induction (Fig. 2F,G). Interestingly, we observe that FOXO3.A3 expression did not induce p53 or p21 expression and induces APC/C^CDH1^ activation in S/G2 of RPE-FUCCI-FOXO3 cells knockout for p53 (Δp53) or p21 (Δp21) (Fig. 2H & S3A-C).

Collectively, these observations imply that FOXOs promote premature APC/C^CDH1^-activation in response to replication stress. In order to test whether FOXOs play an essential role in this process, we generated RPE-FUCCI cells with a doxycycline-inducible shRNA that efficiently knocks down *FOXO1* and *FOXO3* (RPE-FUCCI-shFOXOs) (30–32). Aditionally, to separate any FOXO-dependent effect from p53- and p21-mediated effects, we also introduced the inducible shFOXOs into Δp21 and Δp53 RPE-FUCCI cells. Replication stress induced the expression of p21 in a p53-dependent manner and p53 and p21 expression are reduced in the absence of FOXOs, both in unperturbed conditions and after replication stress (Fig. 3A). While FOXO-dependent p53 activation is not required for replication stress-induced cell cycle exit (Fig. 1G). However, FOXO depletion itself did prevent replication stress-induced premature APC/C^CDH1^ activation in G2, uncovering an additional role for FOXO in the cellular response to replication stress (Fig. 3B,C). Loss of FOXOs lowered the activity of CHK1 and CHK2 in response to aphidicolin and other DNA damaging agents like hydroxyurea (HU) and neocarzinostatin (NCS) (Fig. 3D), but did not prevent initial DNA damage signaling, as FOXO-depletion did not significantly reduce the formation of γH2AX foci in response to aphidicolin treatment (Fig. 3E). Downstream events, such as the loading of mono-ubiquitinated FANCD2 and the establishment of 53BP1 foci in S/G2 cells, following replication stress were perturbed in the absence of FOXO activity (Fig. 3F-H). Together these results delineate an important role for FOXOs in both replication stress checkpoint activation and steering cell fate by stimulating a cell cycle exit following replication stress. This suggests that reducing FOXO expression might increase resistance to aphidicolin treatment. Indeed, loss of FOXOs increased the IC50 of aphidicolin in RPE-FUCCI cells by ~54% (from 113.1nM in control to 174.1nM in FOXO-depleted RPE-FUCCI cells)(Fig. 3I). Taken together, we uncover FOXOs as novel important players in the replication stress response but still wonder what is the event downstream of FOXOs nuclear translocation that drives cell cycle withdrawal. Our data show that FOXOs can drive APC/C^CDH1^ activation following replication stress. Indeed, we observed that the FOXO3.A3-induced loss of Geminin expression in RPE-FUCCI cells (4n mKO+ cells) is reduced upon CDH1 knock-down (Fig. S4A-C). In addition, FOXO3.A3 expression results in a gradual decrease in EMI1 expression in asynchronously cultured cells, which is in line with previous studies showing that EMI1 downregulation is a prerequisite for cell cycle exit in G2 phase (Fig. 4A & Fig. S4D)(9,11,17). To exclude that EMI1 downregulation is not simply the effect of a FOXO-induced G1 arrest, we sorted RPE-FUCCI-FOXO3.A3 cells based on mKO and mAG expression 8 and 16 hours after FOXO3 induction. As expected, EMI1 expression is not detected in G0/G1-phase and high in S/G2-phases in control cells (Fig. 4B,C). Strikingly, FOXO3.A3 induction strongly diminished EMI1 expression in S/G2-phase cells both at the level of mRNA and protein (Fig. 4B-C). Accordingly, RPE-FUCCI cells were more prone to exit the cell cycle when EMI1 knockdown was combined with FOXO3.A3 expression (Fig. 4D & S4E). To determine whether EMI1 expression can prevent FOXO3-induced premature APC/C^CDH1^ activation, we constructed RPE-FUCCI cells with doxycycline-inducible FOXO3.A3 and mTurq2-EMI1 (RPE-FUCCI iFOXO3.A3/mTurq2-EMI1) (Fig. 4E). While FOXO3.A3 expression reduced mitotic entry of G2 cells due to premature APC/C^CDH1^ activation (Fig. 4F), simultaneous expression of EMI1 blocked APC/C^CDH1^ activation in G2, allowing cells to enter mitosis (Fig. 4F). Subsequently, cells arrested at metaphase as a consequence of high EMI1 levels in mitosis (Fig. 4F & S4F)(33).

**Figure 3:**
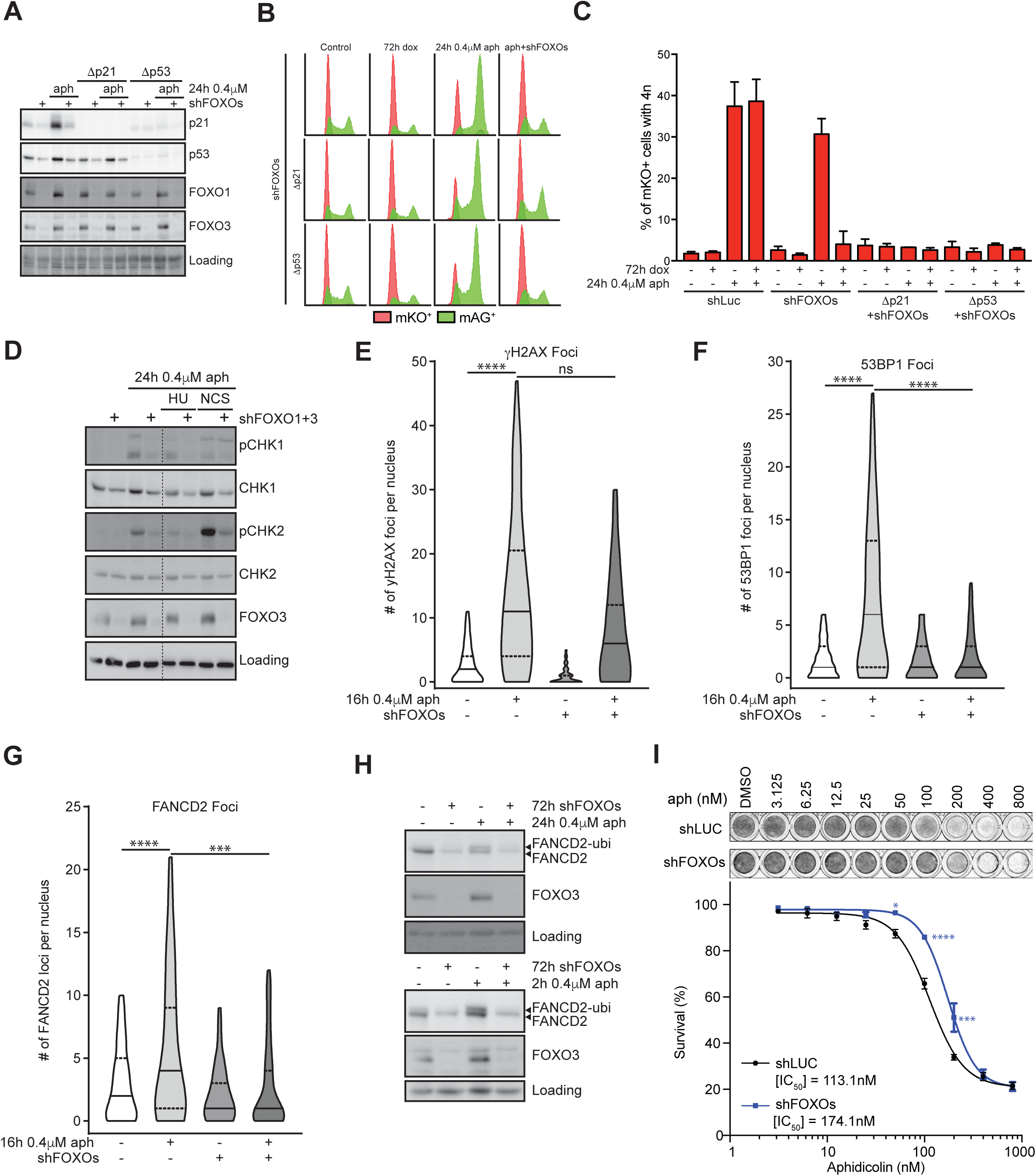
FOXOs support replication stress response and cooperate with p53/p21 to induce cell cycle exit. **A** Westernblot for p21, p53, FOXO1and FOXO3 levels in RPE-FUCCI, p21 (Δp21) and p53 (Δp53) knockout cells before and after 0,4μM aphidicolin treatment in combination with shFOXOs. Nonspecific background staining was used as loading. **B** Representative flow cytometric analysis of hoechst stained RPE-FUCCI, p21 (Δp21) and p53 (Δp53) knockout cells before and after 0,4μM aphidicolin treatment in combination with shFOXOs. Red: CDT1-mKO+ (G1-phase), green: Geminin-mAG+ (S/G2/M-phase). **C** Flow cytometric quantification of 4n CDT1-mKO+ cells after 0,4μM aphidicolin treatment in combination with shFOXOs of p53 (Δp53) and p21 (Δp21) knockout RPE-FUCCI cells (n=3). **D** Westernblot for phospho-CHK1, CHK1, phospho-CHK2, CHK2 and FOXO3 expression levels after 0,4μM aphidicolin treatment in combination with Hydroxyurea (HU) or Neocarcinostatin treatment (NCS). Nonspecific background staining was used as loading. **E** Violin plot representing immunofluorescence quantification of the amount of γH2AX foci in control (n=821), Aph (n=88), shFOXOs (n=191) and combined Aph + shFOXOs (n=71) treated cells. Median = solid line, quartiles = dashed line. Welch’s corrected unpaired t-test P<0.00005=****. **F** Violin plot representing immunofluorescence quantification of the amount of 53BP1 foci in control (n=1553), Aph (n=1059), shFOXOs (n=801) and combined Aph + shFOXOs treated cells (n=1639). Median = solid line, quartiles = dashed line. Welch’s corrected unpaired t-test P<0.00005=****. **G** Violin plot representing immunofluorescence quantification of the amount of FANCD2 foci in control (n=821), Aph (n=273), shFOXOs (n=116) and combined Aph + shFOXOs (n=655) treated cells. Median = solid line, quartiles = dashed line. Welch’s corrected unpaired t-test P<0.0005=***, p<0.00005=****. **H** Westernblot for mono-ubiquitinated FANCD2 and FOXO3 expression levels after 0,4μM aphidicolin treatment for 2 and 24 hours in combination with FOXO knockdown. Nonspecific background staining was used as loading control. **I** Dose response curve of cell viability after treatment with aphidicolin for 7 days in RPE-FUCCI cells with expressing shLuc or shFOXOs. Bonferonni Muiltiple testing corrected ANOVA p-value <0.05= *, p<0.005=***, <0.0005 = ****.

**Figure 4:**
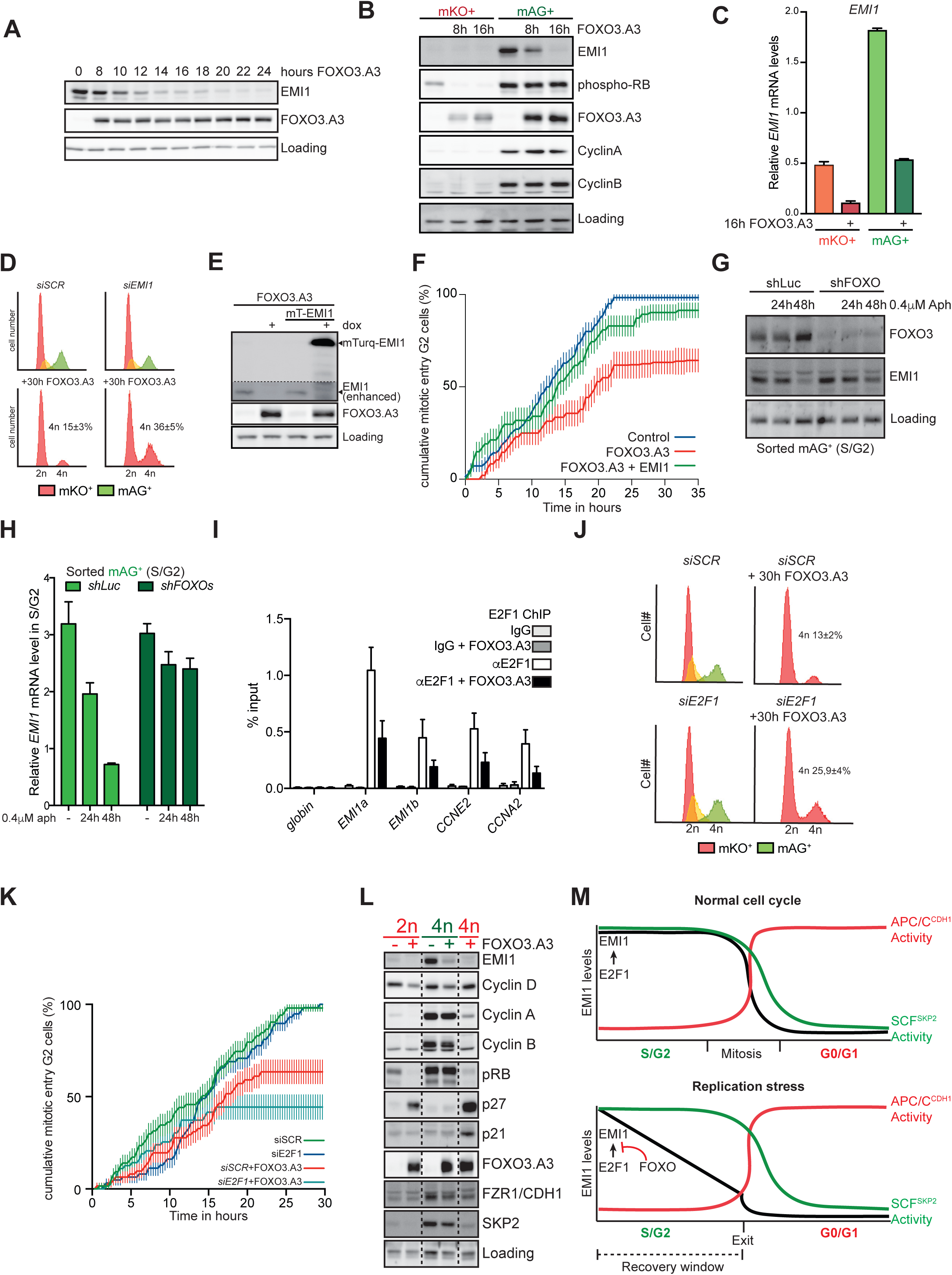
FOXOs activate APC^CDH1^ in G2 by downregulating EMI1. **A** Westernblot for EMI1 and FOXO3 levels after 8, 10, 12, 14, 16, 18, 20, 22 and 24h FOXO activation in asynchronous RPE-FUCCI cells. Nonspecific background staining was used as loading. **B** Westernblot for EMI1, phospho-RB, FOXO3, Cyclin A and Cyclin B in CDT1-mKO+ and Geminin-mAG+ sorted cells after 8 and 16h FOXO activation. Nonspecific background staining was used as loading. **C** qPCR analysis of *EMI1* expression in CDT1-mKO+ and Geminin-mAG+ sorted cells after 16h FOXO activation. **D** Flow cytometric analysis of hoechst stained RPE-FUCCI-iFOXO3.A3 before and after 30h FOXO3.A3 expression in combination with *siEMI1*. % 4n n=3. Red: CDT1-mKO+, green: Geminin-mAG+. **E** Westernblot for EMI1, mTurq2-EMI1 and FOXO3.A3 expression in RPE-FUCCI-iFOXO3.A3+EMI1 cells. Nonspecific background staining was used as loading control. **F** Cumulative mitotic entry of G2 cells that progressed through S phase after FOXO3.A3 and EMI1 overexpression. Average of three independent experiments + sem. **G** Westernblot for EMI1 and FOXO3.A3 expression in mAG+ sorted RPE-FUCCI-shFOXOs cells after 24 and 48h 0,4μM aphidicolin treatment. Nonspecific background staining was used as loading control. **H** qPCR analysis of *EMI1* expression in mAG+ sorted cells after 24 and 48h FOXO activation. **I** Average % input of qPCR of genomic E2F1 binding sites in 2 promoter regions of *EMI1* (EMI1a & Emi1b), *CCNE2* and *CCNA2* after E2F1 chromatin immunoprecipitation in the presence/absence of FOXO3.A3 expression (n=3). **J** Flow cytometric analysis of hoechst stained RPE-FUCCI-iFOXO3.A3 before and after 30h FOXO activation in combination with siE2F1. % 4n n=3 **K** Cumulative mitotic entry of G2 cells that progressed through S phase after FOXO3.A3 overexpression in the precence or absence of *siSCR* and *siE2F1*. Average of three independent experiments + sem. **L** Westernblot for EMI1, Cyclin B, Cyclin A, Cyclin D, phospho-RB, p27, p21, FOXO3, CDH1 and SKP2 levels in 2n mKO+, 4n mAG+ 4n mkO+ sorted RPE-FUCCI-iFOXO3.A3 cells after 24 hours FOXO activation. Nonspecific background staining was used as loading. **M** Schematic illustrating tthat FOXOs reduce the levels of EMI1 in response to replication stress. When EMI1 levels reach critically low levels, the APC/C^CDH1^ (red line) activates, SCF^SKP2^ (green line) subsequently is inactivated and cell cycle exit from G2 is triggered.

We noticed that endogenous EMI1 levels still decreased in FOXO3.A3- and EMI1-overexpressing cells (Fig. 4E), suggesting that transcriptional repression by FOXOs is responsible for reduced endogenous EMI1 expression. If FOXOs regulate EMI1 levels in S/G2 phase in response to replication stress, loss of FOXOs should stabilize EMI1 levels. Indeed, aphidicolin-induced loss of EMI1 in S/G2-phase cells is prevented upon FOXO depletion (Fig. 4G-H). Hence, we conclude that FOXO-induced APC/C^CDH1^-activation is mediated via repression of *EMI1*.

Although ~20% of all genes in the genome have FOXO binding elements in their promoters, FOXO binding sites are absent directly upstream of *EMI1*, implying that *EMI1* transcription is not suppressed by FOXOs binding to its promoter directly (34,35). The main transcription factor driving *EMI1* expression is E2F1 (21), and previous studies have shown that FOXOs can bind to E2F1 and alter E2F1 target gene expression (32). Indeed, co-immunoprecipitation (co-IP) experiments using GFP-FOXO1/3 and HA-E2F1 confirmed that FOXOs can bind to E2F1 (Fig. S5A). To test whether FOXOs indeed affect E2F1-dependent transcription of *EMI1* and other genes, we performed Chromatin-IP (ChIP) experiments. We found that binding of endogenous E2F1 to the promoter of *EMI1* as well as two other canonical E2F1 targets (Cyclin E (*CCNE2*) and A (*CCNA2*)) was reduced when FOXO3.A3 was expressed (Fig. 4I). Accordingly, the expression of E2F1 target genes (*E2F1* itself, *CCNE2* and *CCNA2*) was reduced (Fig. S5B-D). Indeed, FOXO-induced APC/C^CDH1^ activation is enhanced when cells enter S-phase with lowered E2F1 levels, as combining *E2F1* knockdown with FOXO3.A3 expression sensitizes RPE-FUCCI cells to activate APC^CDH1^ in S/G2 (Fig. 4J,K & S6E).

Next, we determined whether cells that activate APC/C^CDH1^ in G2-phase truly switch to a G0/G1 state after EMI1 downregulation. To this end, we sorted RPE-FUCCI-FOXO3.A3 cells based on mAG, mKO and DNA content (Hoechst 33342) at 24 hours after FOXO3.A3 induction (Fig. 4L). 2n mKO+ cells reflect G1 as EMI1 levels and RB phosphorylation are low, Cyclin D is high and both Cyclin A and B are absent. FOXO3.A3 expression in 2n mKO+ cells induced expression of its target p27 and reduced the expression of Cyclin D, known to lead to a G1 arrest (29). In 4n mAG+ control cells the expression of EMI1, Cyclin A, Cyclin B, SKP2, CDH1 and phosphorylation of RB are high, confirming that these cells are indeed in S/G2-phase. Strikingly, FOXO3 activation reduced EMI1 expression in 4n mAG+ cells, illustrating that FOXO mediated EMI1 repression precedes APC/C^CDH1^ activation. Additionally, P27 and p21 are not induced, in line with the fact that SCF^SKP2^ mediates their degradation in S/G2 cells (36,37). Importantly, 4n mKO+ cells resemble G1 cells as expression of EMI1, Cyclin A, Cyclin B, SKP2 and phosphorylation of RB are absent. The high p27 and p21 levels in 4n mKO+ cells suggest that these cells are arrested. Finally, this experiment also confirms our initial observation that APC^CDH1^ activation is not preceded by CDK inhibition as p27 and p21 only increase after the APC/C is activated. Collectively these data show that FOXO-dependent premature activation of the APC/C^CDH1^ prevents mitotic entry and in stead reverts the cells back into a 4N G1-like cellular state.

## Conclusion & Discussion

Here we investigated the cellular response to replication stress, and find that cells monitor the presence of residual damage when progressing from S to G2 phase. We identified that a fraction of cells initiates cell cycle exit from G2 phase in response to replication stress. This is in contrast to earlier work that suggested that both transformed and non-transformed cells that experienced replication stress progress through mitosis, resulting in chromosomal aberrations and 53BP1-foci in G1-phase (3–5). In agreement with previous reports, we show that G2 phase can be extended in an ATR-dependent manner, to provide time for DNA repair (5). However, if repair is not successful within ~24 hours, APC/C^CDH1^ is activated and cells exit the cell cycle. It has previously been shown that DSBs in G2 phase may cause premature APC/C^CDH1^-activation, resulting in irreversible withdrawal from the cell cycle (9–12,17). Premature APC/C^CDH1^-activation is preceded by p21-dependent nuclear entrapment of Cyclin B1-CDK complexes, which renders it inert for CDK (re-)activation (10,18). Intriguingly, here we find that CDK2 remains active during a replication stress-induced G2 arrest, and that APC/C^CDH1^ can be activated independent from p53, p21 or CDK2 inhibition. This leads to the interesting conclusion that APC/C^CDH1^ can be activated after replication stress in cells containing active CDK-complexes. We find that aphidicolin-induced APC/C^CDH1^-activation is dependent on the repression of EMI1. FOXOs act as a timer to restrict the window during which cells may recover from replication stress, by gradually decreasing EMI1 levels through removing E2F1 from the *EMI1* promoter (Fig. 4M). Alteration of E2F1 dependent transcriptional output by FOXOs has been reported previously for genes involved in apoptosis, and we now show that this mechanism of action is also applicable to cell cycle regulation in S/G2-phase (32).

Interestingly, FOXOs and p53/p21 co-operate to restrict the time during which replication stress induces a reversible cell cycle arrest. On the one hand, FOXOs support the checkpoint and sensitize the APC/C^CDH1^ to activation, on the other hand p21 inhibits CDK activity. Combined, this results in a robust switch in which APC/C^CDH1^-activation can revert the cell from a S/G2 state into G0/G1. Next to indirectly regulating the APC/C^CDH1^, we observed that FOXOs are required for the establishment of the replication stress checkpoint. As FOXOs have been implicated to play a similar role in ATM/CHK2 mediated DSB repair, it will be interesting to investigate the FOXO-dependent mechanisms controlling the establishment of the replication stress checkpoint (38–40).

Noteworthy, FOXOs have been suggested to play a role in inducing a G2 arrest, but these conclusions were based solely on DNA content analysis (41–44). As we do not observe mAG+ cells after FOXO activation, it’s tempting to speculate that the previously reported FOXO-induced G2 arrest in fact represent cells that exited G2 and are in a G0/G1 state with 4n DNA.

Combined, our results establish a novel role for FOXOs in the replication stress response. FOXOs cooperate with the ATR/CHK1 and p53/p21 response both by supporting the establishment of a checkpoint and simultaneously restricting the time in which cells are allowed to resolve the damaged DNA. Importantly, deregulation of this FOXO driven timer gives cells the opportunity to divide with higher levels of replication stress, potentially promoting cellular transformation.

## Materials & Methods

### Cell culture

hTERT-immortalized Retinal Pigment Epithelial cells (RPE-1) were cultured in DMEM-F12 (Lonza) containing 10% FBS (Bodinco), 100U/ml penicillin and 100mg/ml streptomycin (Lonza). For doxycycline treatment 200 ng/ml Doxycycline (Sigma) was used for the duration of indicated times. RPE-1 cells in which a fluorescent tag was introduced in one allele of Cyclin B1 (RPE CCNB1^YFP^) have been described before (43). RPE-CCNB1^YFP^-53BP1^mCherry^, RPE CCNB1^YFP^-EMI1^mTurq2^-53BP1^mCherry^ cells and RPE-CCNB1^YFP^-DHB^mCherry^ cells were described before (11). Generation of RPE-FUCCI cells was previously described. Cells were treated with 0,2μM, 0,4μM aphidicolin (Sigma) for the indicated times. 0,2μM is used to induce replication stress from which cells can recover. 0,4μM induces a S/G2 arrest and was used to study APC^CDH1^ activation and replication stress checkpoint establishment.

### Constructs, lentiviral transduction and transfections

Lentiviral cDNA expression vectors expressing FOXO3.A3 were generated using Gateway cloning in the pINDUCER20 (Addgene #44012) doxycycline inducible expression system (45). The lentiviral construct pCW-mTurqouise-EMI1 was generated as previously described (19). Transfecting third generation packaging vectors using Poly-ethylenimine into HEK293T cells generated lentiviral particles. Transfection of pTON-BIOPS-Flag-EGFP-FOXO1/FOXO3 and pCDNA-HA-E2F1 in HEK293T cells was performed using ExtremeGene 9 (Roche). Transfection of siRNA smartpools targeting *EMI1*, *FZR1/CDH1* and *E2F1* was performed using lipofectamine (Life Technologies).

CRISPR/Cas9 mediated knockouts we generated as previously described in (46) Briefly, pX330 was transfected in for both TP53 and CDKN1A knockout generation. Subsequently, cells were treated with 5μM Nutlin-3a for 7 days to select out TP53 and CDKN1A deficient cells.

### Immunoblotting & antibodies

For western blot cells are lysed, protein concentration is measured (Bradford, BioRad) and finally dissolved in sample buffer containing 0.2%SDS, 10% glycerol, 0.2% β-mercapto-ethanol, 60mM Tris pH6.8. Equal protein concentrations were loaded and proteins were detected using 6-15% SDS-PAGE gels and subsequent western-blot analysis with primary antibodies recognizing FOXO1 (CS2880), FOXO3 (NB100614), p27 (BD-610241), p21 (BD556430), p53 (SC-126), EMI1 (ThermoFisher 3D2D6), E2F1 (CST-3742), RB-S807/811 (CST-9308), Cyclin A (AB-16726), Cyclin E (SC-198), Cyclin D (ab134175), Cyclin B (SC-752), CDH1/FZR1 (MS-1116-P1), SKP2 (CST-2652S), CHK1 (SC8408), phospho-CHK1 (CD2348), CHK2 (SC9064), phospho-CHK2 (CS2261), FANCD2 (ab108928), 53BP1 (NB100-304), yH2AX (MP 05-636), HA (home made), GFP (home made), used 1:2000. Primary antibodies were detected by secondary HRP conjugated antibodies targeting mouse, rabbit, and rat IgG and visualized using chemiluminescence (Biorad) and an ImageQuant LAS 4000 scanner (GE Healthcare). For immunoprecipitation cells transfected with HA-E2F1, GFP-FOXO1, GFP-FOXO3 were lysed in 50mM Tris pH7.4, 150mM NaCl, 1% Triton-X100. For immunoprecipitation Chromotec GFP beads were incubated with cell lysate for 2 hours at 4°C, subsequently washed with lysis buffer and boiled in sample buffer. Chromatin-IP was performed as previously described using rabbit anti E2F1 (C-20, SC193) and Rabbit IgG (SC2027) (34)

### Immunofluorescence

Cells were grown on glass coverslips, fixed using 4% paraformaldehyde and blocked with PBS containing 2% bovine serum albumin (BSA) (Invitrogen) and 0.1% normal goat serum (Invitrogen). Cells were incubated with indicated antibodies (1:500), secondary Alexa488/561 conjugated antibodies and Hoechst (Sigma). Slides were imaged on a Zeiss LSM710 confocal microscope. Quantification of FOXO localization and 53BP1, FANCD2 and γH2AX foci was performed with a custom written ImageJ script that was described previously (19),

### RT-qPCR

mRNA was isolated from live cells using the Qiagen RNeasy kit (Qiagen) and cDNA synthesis was performed using the iScript cDNA synthesis kit (BioRad). Real-time PCR was performed using Fast Start Universal SYBR Green Master (ROX) mix (Roche) in the CFX Connect Real-time PCR detection system (BioRad). Target genes were amplified using specific primer pairs (Supplemental table 1) and specificity was confirmed by analysis of the melting curves. Target gene expression levels were normalized to *GAPDH* and *HNRNPA1* levels.

### Flow cytometry

For DNA content profiling and sorting, live cells were incubated with 10 μg/ml Hoechst33342 for 30 min. at 37°C. After incubation cells were trypsinized and transferred to normal culture medium before measuring. mKO-hCDT1, mAG-hGeminin and Hoechst33342 intensity was measured using a BD LSR Fortessa Flow cytometer or BD Aria II FACS (BD bioscience).

### Live cell imaging and tracking

20.000 RPE cells were cultured in Lab-Tek II 8-well imaging chambers. Prior to imaging normal culture medium is replaced with Leibovitz medium (Lonza) containing 10% FCS (Lonza),2 mM L-Glutamin, 100U/ml penicillin and 100μg/ml streptomycin (Lonza). Cells were treated with 0.4 μM aphidicolin and/or doxycycline (Sigma) at indicated timepoints before imaging. For UV irradiation, medium was aspirated and cells were rinsed with PBS before exposure to global UV irradiation by TUV lamp. After irradiation Leibovitz’s L-15 (Gibco) CO2-independent medium, supplemented with ultra-glutamine, penicillin/streptomycin and 10% fetal calf serum was added to start live-cell imaging as described above. For all experiments where phenotypic outcome was quantified at least 50 cells per condition in each independent biological replicate were scored, n≥50, unless otherwise stated in figure legends. Imaging was performed on Zeiss Cell observer Real-Time imaging and DeltaVision Elite (applied precision) microscopes for 48 hours at 37°C. Cell tracking and quantification was performed using ImageJ. Cells expressing mAG-hGeminin at the moment of doxycycline addition and cell starting to express mAG-hGeminin within 3 hours after doxycycline addition were considered S/G2 in the analysis. Relative CDK2 activity and cyclin B1^YFP^ intensity were measured in individual G2 cells that degraded Cyclin B1 after progression through S phase in presence of 0,4µM aphidicolin. Cells were in silico alligned at the onset of cyclin B1 degradation and cyclin B1^YFP^ values were normalized to the max cyclin B1 level.

### mRNA sequencing & Analysis

RPE-1 hTERT cells were synchronized with a double-thymidine block and released in the absence (DMSO) and presence of aphidicolin (DRS) and cultured for 7 hours. Subsequently, total RNA from cultured cells was extracted using TRIzol reagent (Invitrogen). Strand-specific libraries were generated using the TruSeq PolyA Stranded mRNA sample preparation kit (iIlumina). In brief, polyadenylated RNA was purified using oligo-dT beads. Following purification, the RNA was fragmented, random-primed and reserve transcribed using SuperScript II Reverse Transcriptase (Invitrogen). The generated cDNA was 3′ end-adenylated and ligated to Illumina Paired-end sequencing adapters and amplified by PCR using HiSeq SR Cluster Kit v4 cBot (Illumina). Libraries were analyzed on a 2100 Bioanalyzer (Agilent) and subsequently sequenced on a HiSeq2000 (Illumina). We performed RNaseq alignment using TopHat 2.1.1 (Supplemental table 2) (47). Differentially expressed genes were called with DEseq2 (48) with an adjusted p value threshold of <0,005. Transcription factor binding site analysis was performed using oPOSSUM (49) to determine enrichment for transcription factors by Z-score and Fisher exact score over the whole genome compared to our upregulated genes (Supplemental table 2).

## Author contributions

MH, FF, LK, LS, JB, BB performed experiments. MH, FF, LK, JB wrote the manuscript. NN, MR supported cell sorting. TD, RM, BB financially supported this study and provided critical feedback.

## Acknowledgements

This study is financially supported by CancerGenomicsCenter.nl, the Oncode institute and the research grants UU2014-6902, UU2009-4490 (to TD) and NKI2017-6787 (to RM) from the Dutch Cancer Society (KWF kankerbestrijding).

---

**Supplementary figure 1.**
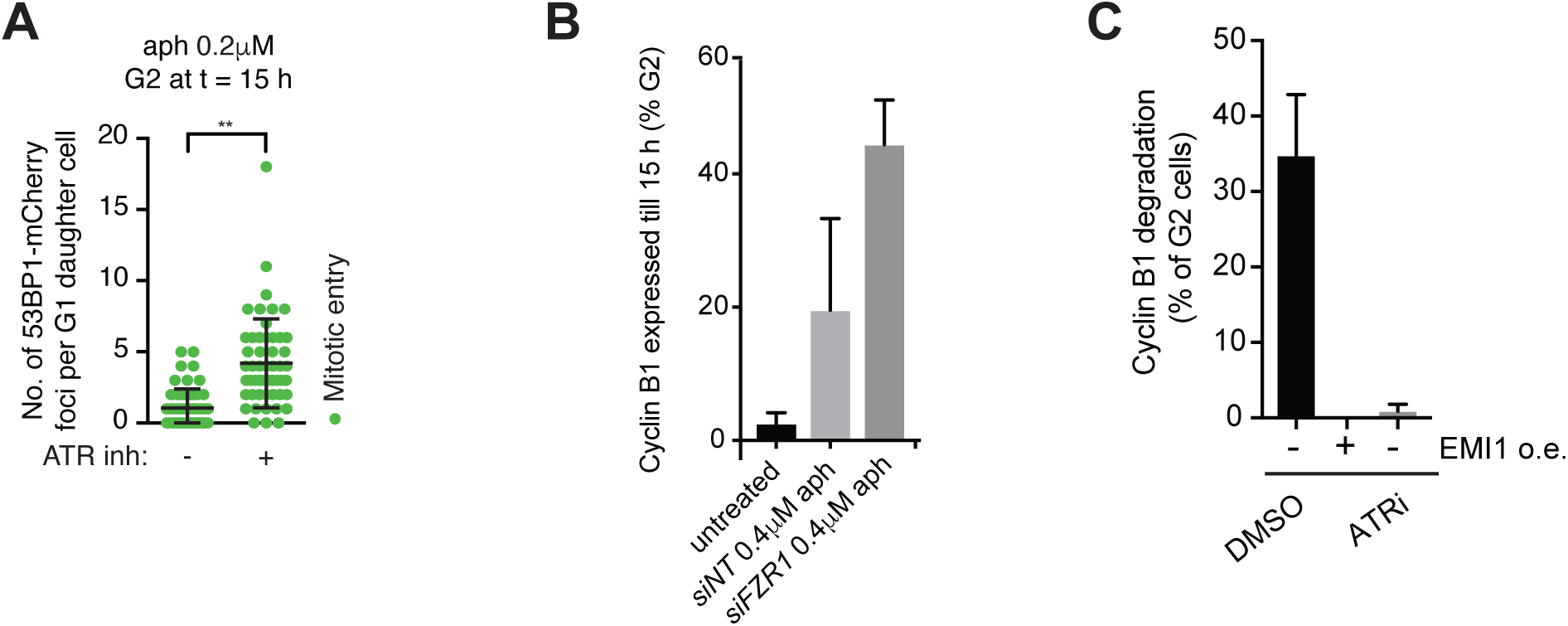
Quantification of 53BP1mCherry foci in G1 after mitotis in the precense of 0,2mM aphidicolin and/or ATR inhibitor for >15h. Dots represent individual cells pooled from two independent experiments. Mean + sd. ** P<0.001 (unpaired t-test). **B** Percentage of G2 cells that retain cytoplasmic cyclin B1 expression during the course of the exp (16 h) following aphidicolin treatment in presence of indicated knock-down. **C** Percent of G2 cells that degrade Cyclin B1 after combinational treatment of 0,4μM aphidicolin with mTurq2-EMI1 expression (dox) or 5μM ATR inhibitor VE-821.

**Supplemental figure 2.**
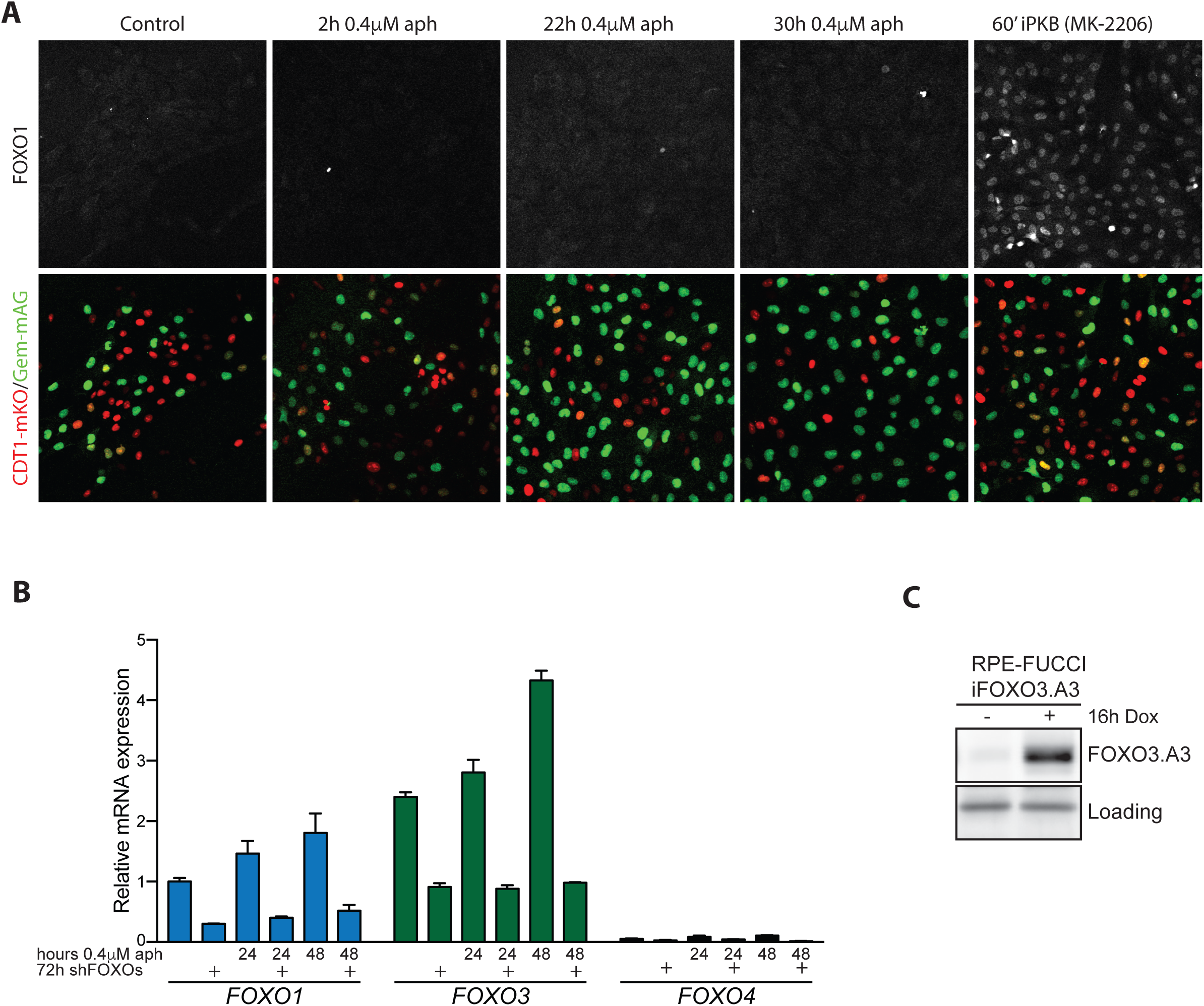
**A** Immunofluorescence image of FOXO1 (Grey) expression and localization in RPE-FUCCI cells (CDT1 (G1) = Red, Geminin (S/G2/M) = green) after Aph or 1μM AKT inhibitor treatment. **B** RT-qPCR analysis of *FOXO1, FOXO3* and *FOXO4* mRNA expression level changes in RPE-FUCCI-shFOXOs cells during 48 hours of 0,4μM Aph treatment in the absence or presence of shFOXOs. **C** Westernblot for FOXO3.A3 expression after 16h doxycycline (dox) treatment. Nonspecific background staining was used as loading.

**Supplemental Figure 3.**
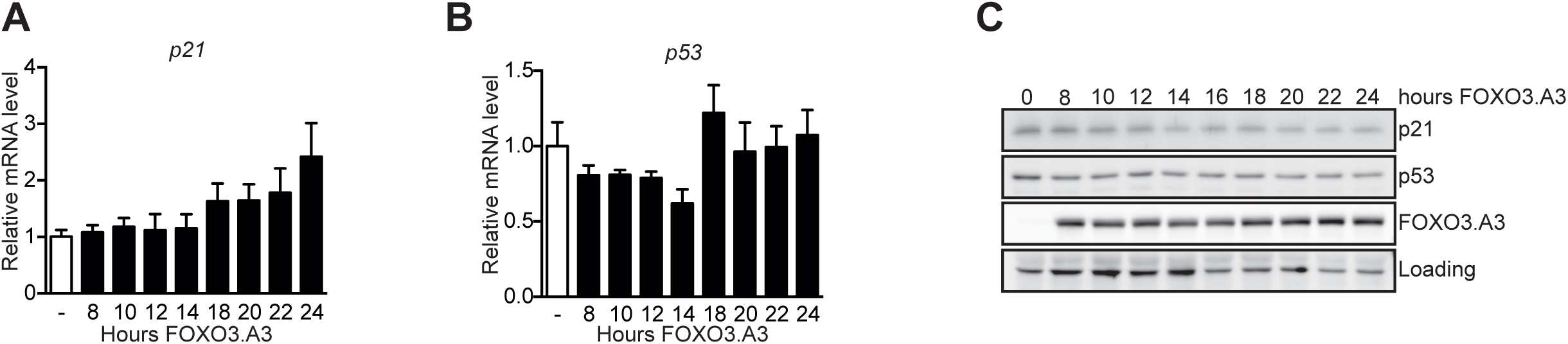
**A&B** RT-qPCR analysis of *P21* and *P53* mRNA expression level changes in RPE-FUCCI-iFOXO3.A3 cells over the course of 24 hours doxycycline (dox) treatment. **C** Western blot analysis of p21, p53 and FOXO3 protein levels in RPE-FUCCI-iFOXO3.A3 after doxycycline treatment for 24 hours. Equal concentrations of protein are loaded and aspecific background staining is used to visualize equal loading.

**Supplemental figure 4.**
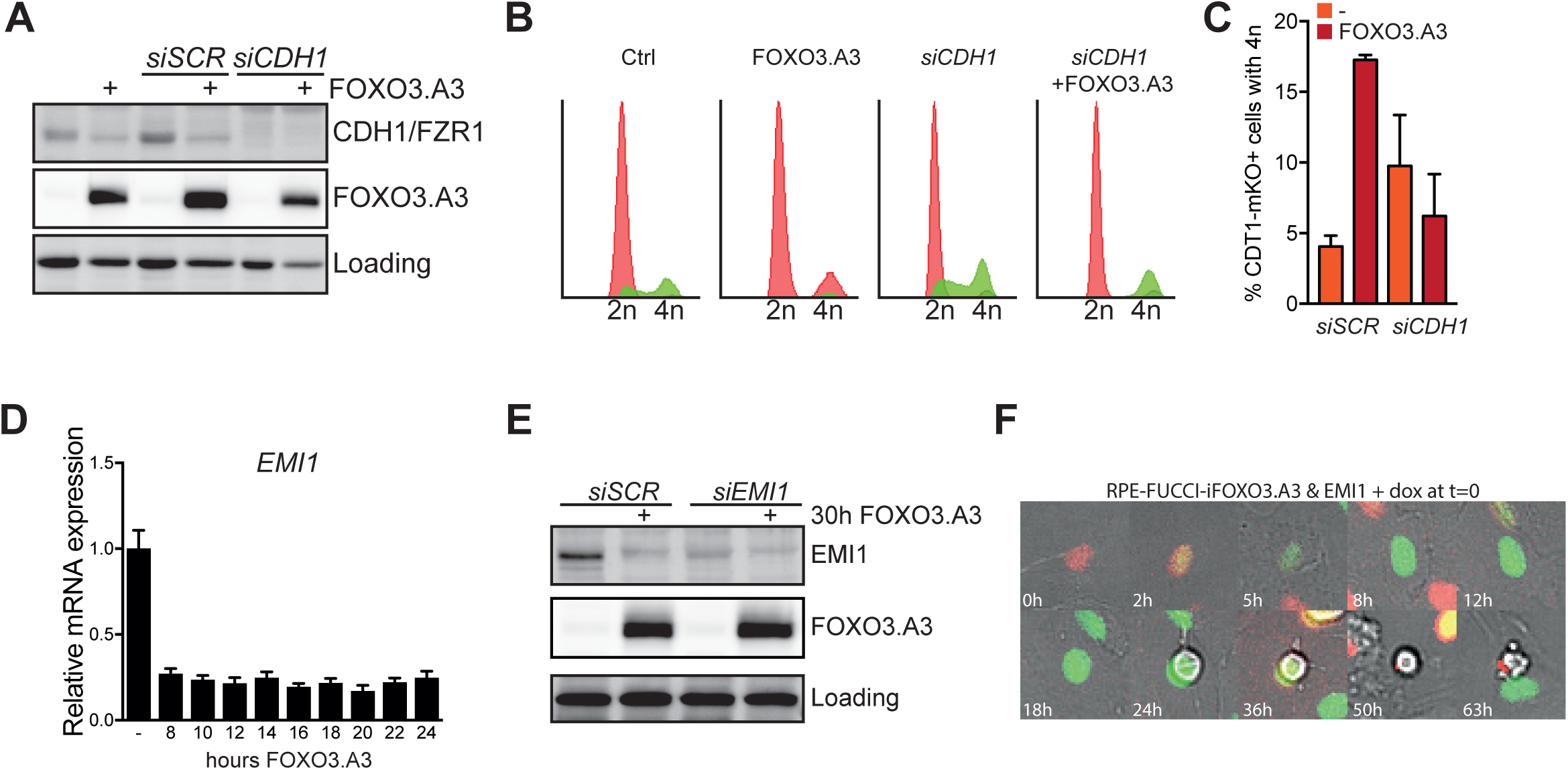
**A** Western blot analysis of FZR1/CDH1 protein levels in RPE-FUCCI-iFOXO3.A3 after 72 hours of siSCR or siCDH1 with doxycycline treatment for 30 hours. Equal concentrations of protein are loaded and aspecific background staining is used to visualize loading. **B** Representative flow cytometric analysis of hoechst stained RPE-FUCCI-iFOXO3.A3 before and after 30h FOXO activation in combination with siCDH1. Red: CDT1-mKO+ (G1-phase), green: Geminin-mAG+ (S/G2/M-phase). **C** Flow cytometric quantification of 4n G1 cells after 30h FOXO activation in combination with siCDH1 (n=3). **D** RT-qPCR analysis of *EMI1* expression in RPE-FUCCI-iFOXO3.A3 cells over the course of 24 hours FOXO3.A3 expression. **E** Western blot analysis of EMI1 protein levels in RPE-FUCCI-iFOXO3.A3 after 72 hours of siSCR or siEMI1 in combination with doxycycline treatment for 30 hours. Equal concentrations of protein are loaded and aspecific background staining is used to visualize loading. **F** Representative live imaging sequence of RPE-FUCCI-FOXO3.A3+EMI1 cells visualizing the mitotic arrest observed after combined FOXO3 and EMI1 overexpression (dox). Red = mKO2 (G1 phase), Green is mAG. (S/G2/M phase)

**Supplemental figure 5.**
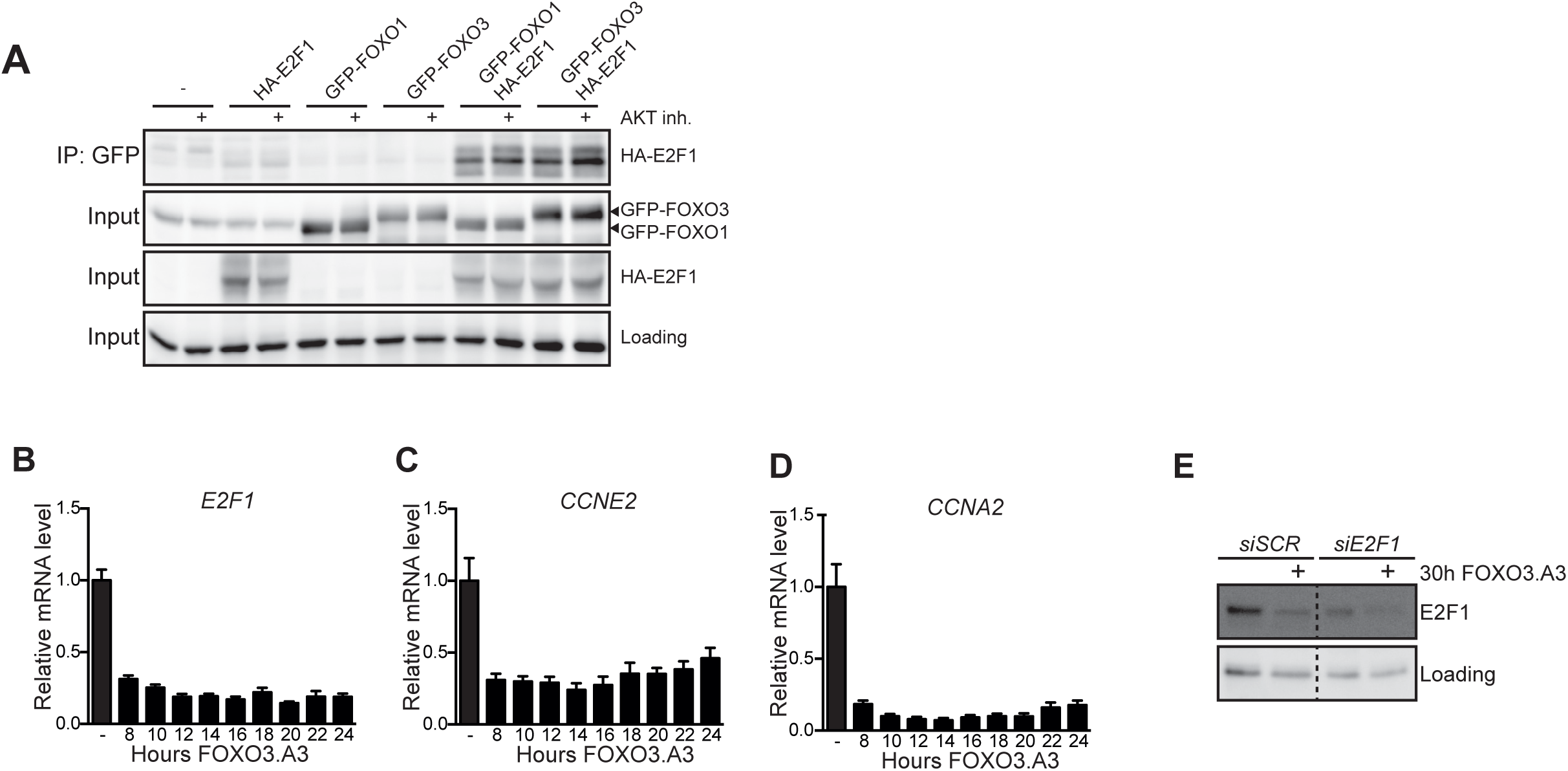
**A** Westernblot for HA-E2F1, GFP-FOXO1 and GFP-FOXO3 protein levels after GFP immunoprecipitation in the presence/absence of 10μM AKT inhibitor VIII for 60 min. Nonspecific background staining was used as loading. **B-D** RT-qPCR analysis of *E2F1, CCNE2, CCNA2* mRNA expression level changes in RPE-FUCCI-iFOXO3.A3 cells over the course of 24 hours doxycycline treatment (FOXO ON). **E** Western blot analysis of E2F1 protein levels in RPE-FUCCI-iFOXO3.A3 after 72 hours of *siSCR* or *siE2F1* in combination with doxycycline treatment for 8 hours. Equal concentrations of protein are loaded and aspecific background staining is used to visualize equal protein content.

